# Exploring Food Contents in Scientific Literature with FoodMine

**DOI:** 10.1101/2019.12.17.880062

**Authors:** Forrest Hooton, Giulia Menichetti, Albert-László Barabási

## Abstract

Thanks to the many chemical and nutritional components it carries, diet critically affects human health. However, the currently available comprehensive databases on food composition cover only a tiny fraction of the total number of chemicals present in our food, focusing on the nutritional components essential for our health. Indeed, thousands of other molecules, many of which have well documented health implications, remain untracked. To explore the body of knowledge available on food composition, we built FoodMine, an algorithm that uses natural language processing to identify papers from PubMed that potentially report on the chemical composition of garlic and cocoa. After extracting from each paper information on the reported quantities of chemicals, we find that the scientific literature carries extensive information on the detailed chemical components of food that is currently not integrated in databases. Finally, we use unsupervised machine learning to create chemical embeddings, finding that the chemicals identified by FoodMine tend to have direct health relevance, reflecting the scientific community’s focus on health-related chemicals in our food.

## Introduction

Decades of research in nutrition have documented the exceptional role of diet in health, unveiling the role of selected nutrients, like sugars, fats, proteins, vitamins, and other biochemical factors, as well as factors contributing to non-communicable diseases like deficiency diseases, cardiovascular disease, obesity, and diabetes mellitus. However, our ability to explore how food affects our health is severely limited by the lack of systematic knowledge on food composition. Among the national databases for food composition, the most accurate data is maintained by USDA, tracking 188 biochemicals, often called nutritional components.^1,2^ Yet, when it comes to the biochemical composition of the food we consume, these nutritional components represent only a tiny fraction of definable biochemicals reported in food. For example, FooDB, a database that integrates food composition data from databases like USDA, Frida, Duke, Phenol Explorer, and others, catalogues altogether 26,625 compounds.^3–6^ The majority of these compounds are only identified, with no information on their quantities in specific ingredients. For example, sulfides are reported to be present in the Allium family, like garlic or onion, but the precise quantities for important sulfides like diallyl disulfide (garlic) and dipropenyl sulfide (onion) remain unknown, despite their well-documented role in cancer prevention.^7–10^ The current incomplete knowledge of the full biochemical composition of food impedes the research community’s ability to uncover the mechanistic effect of the thousands of untracked molecules and their ultimate mechanistic roles in health, achieved either through the microbiome^11^, by contributing to the body’s metabolism, or by regulating molecular processes in human cells.

The lack of centralized information on the chemical composition of food does not equal a lack of scientific or commercial interest in these chemicals: an exceptional amount of research focuses on identifying and quantifying the presence of certain chemicals in various foods, as well as the health implications and the biochemical roles of specific food-borne chemicals. The problem is that data on the chemical composition of food is scattered across the multiple research literatures, spanning different scientific communities, from agriculture to food research, and from health sciences to biochemistry. While we witness notable efforts to partially mine this extensive literature and catalogue the scattered data into databases, like Phenol Explorer’s focus on polyphenols or eBASIS’s prioritization of human intervention studies^6,12,13^, we lack efforts to achieve this across the full food supply and chemicals.

The lack of systematic efforts to map out the existing information on food prompted us to ask how much information is really available on food composition. We developed FoodMine, a pilot project designed to systematically mine the scientific literature to identify and collect all the chemical compositional data for specific ingredients. Hence we demonstrate the capabilities offered by FoodMine by focusing on garlic and cocoa, foods with well documented health effects, which suggests the existence of a sizable yet scattered literature pertaining information on their chemical contents.^14,15^ The knowledge gathered here serves as a pilot towards future comprehensive systematic efforts aimed at identifying and organizing the available information on the chemical composition of all food throughout the whole scientific literature.

## Results

The FoodMine protocol leveraged the PubMed databases to systematically analyze the title and the abstract of the research papers related to garlic and cocoa (Figure 1).^16^ We entered each food as a search term, obtaining 5,676 papers for garlic and 7,620 papers for cocoa. We subset the search results by applying text matching between predefined vocabularies, MeSH terms^17^ and the abstract of the paper listed in the PubMed entry, narrowing the results to 415 papers for garlic and 475 papers for cocoa. After obtaining the subset of results, we manually accessed the paper if we could access a “full text link”, downloading 299 papers for garlic and 324 papers for cocoa. Finally, we manually evaluated each paper to identify relevant chemical contents and extract information from it. Of the 623 manually evaluated papers, 77 papers contained chemical composition data for garlic and 93 for cocoa, yielding 1,426 and 5,855 individual chemical measurements in total for garlic and cocoa, respectively (see Supplementary Material Section 1). In the resulting FoodMine database a compound is “quantified” when chemical measurements report absolute contents, and “unquantified” otherwise. Figure S1 shows that the majority of papers for both foods contained only one or at most a few chemical measurements, referred to as records. However, an outlier for cocoa reported 960 records^18^, measuring the contents of several compounds in cocoa sourced from 15 different origins; the permutations of these variables resulted in the high number of data points. Another outlier examined a large spread of compounds related to human taste perception, reporting 68 unique compounds.^19^ For garlic, the most relevant outlier reported 198 records.^20^

**Figure 1:**
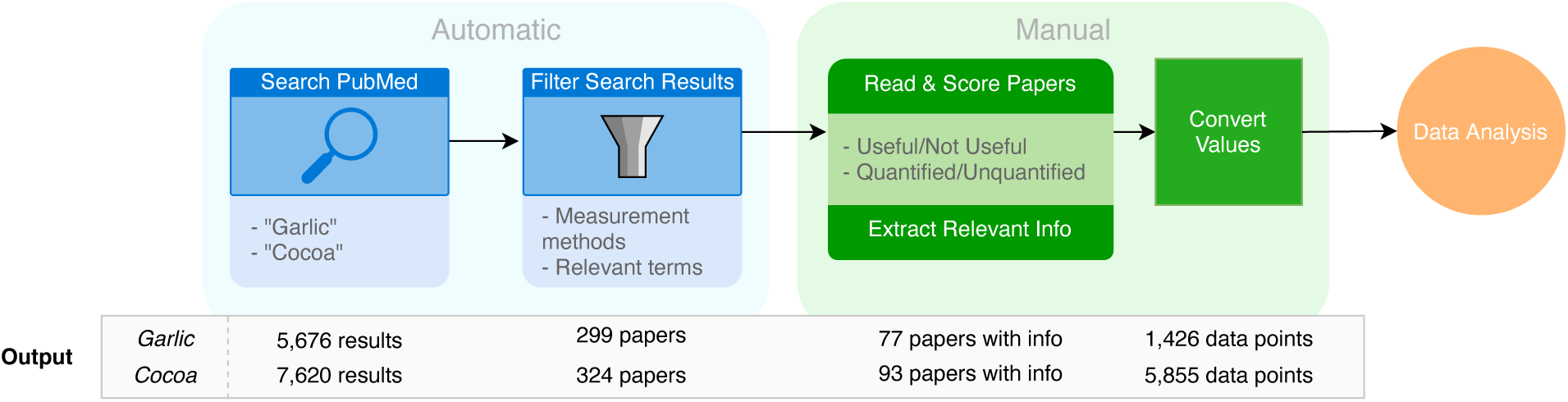
Overview of Data Collection Process. Starting from PubMed, we retrieved a list of paper titles and abstracts using the Pubmed Entrez API, and then applied text matching to automatically filtered the search results, obtaining a subset of papers, which were then read and manually evaluated. If papers contained information on the chemical content of cocoa or garlic, we manually extracted the relevant information. Finally, we converted values in comparable units. The “Output” bar shows the result of each step for garlic and cocoa.

We integrated the compound records into single compound entries, and manually divided quantified entries into their respective compound class based on FooDB classifications, as shown in Figure S2. We find that ‘Carboxylic Acids and Derivatives’ contains the most explored compounds for both garlic and cocoa, and the ‘Flavonoids’ class is in the top three for both ingredients. Compounds from these two classes are common in plant-based food, hence are expected to be present in garlic and cocoa. We also uncovered reports pertaining to various metallic classes, ‘Toxins’, and ‘Pesticides’. Many compounds in the pesticides class came from a paper focusing on the pesticide residues in cocoa products from local markets in Southwestern Nigeria.^21^ Despite its local focus, the examined compounds could directly affect health outcomes worldwide, as Nigeria is the world’s 3^rd^ largest exporter of cocoa.^22^

The FooDB and USDA databases allowed us to verify if the information recovered from the literature matches or contrasts the existing knowledge on the composition of these foods (see Supplementary Material Section 2 for a detailed description of the comparison methodology). To maximize the coverage of this analysis we merged different variations of garlic and cocoa within the USDA and FooDB databases, like merging “Garlic” and “Soft-necked Garlic” in FooDB when comparing the information to FoodMine. In USDA, all reported compounds are quantified, while FooDB lists both quantified and unquantified compounds. We consider a compound quantified if at least one absolute measurement is reported for the selected foods. Taken together, we find that FoodMine recovered more unique compounds than catalogued by USDA (Figure 2A and 2B), and more quantified compounds than catalogued by FooDB. While only 7-9% of compounds are quantified in FooDB and USDA for garlic and cocoa, through FoodMine we collected quantified information for 70% of garlic compounds and 66% of cocoa compounds (see Supplementary Material Section 3). For cocoa and garlic, FooDB and USDA contain more unquantified compounds than quantified. However, we find that ∼70% of the information reported in the literature was quantified, indicating that the literature contains an extensive body of information currently not recorded in databases (see Supplementary Material Section 3). Furthermore, 96 quantified garlic compounds and 283 quantified cocoa compounds are novel, meaning that they were not previously linked to the two ingredients in USDA or FooDB. In summary, 48% and 72% of quantified compounds are novel in both garlic and cocoa, respectively, hence the average increase in quantified measurements offered by FoodMine exceeds 137% (see Supplementary Material Section 3). These findings suggest that a systematic mining of the information scattered in the scientific literature could significantly improve our current knowledge of food composition.

**Figure 2:**
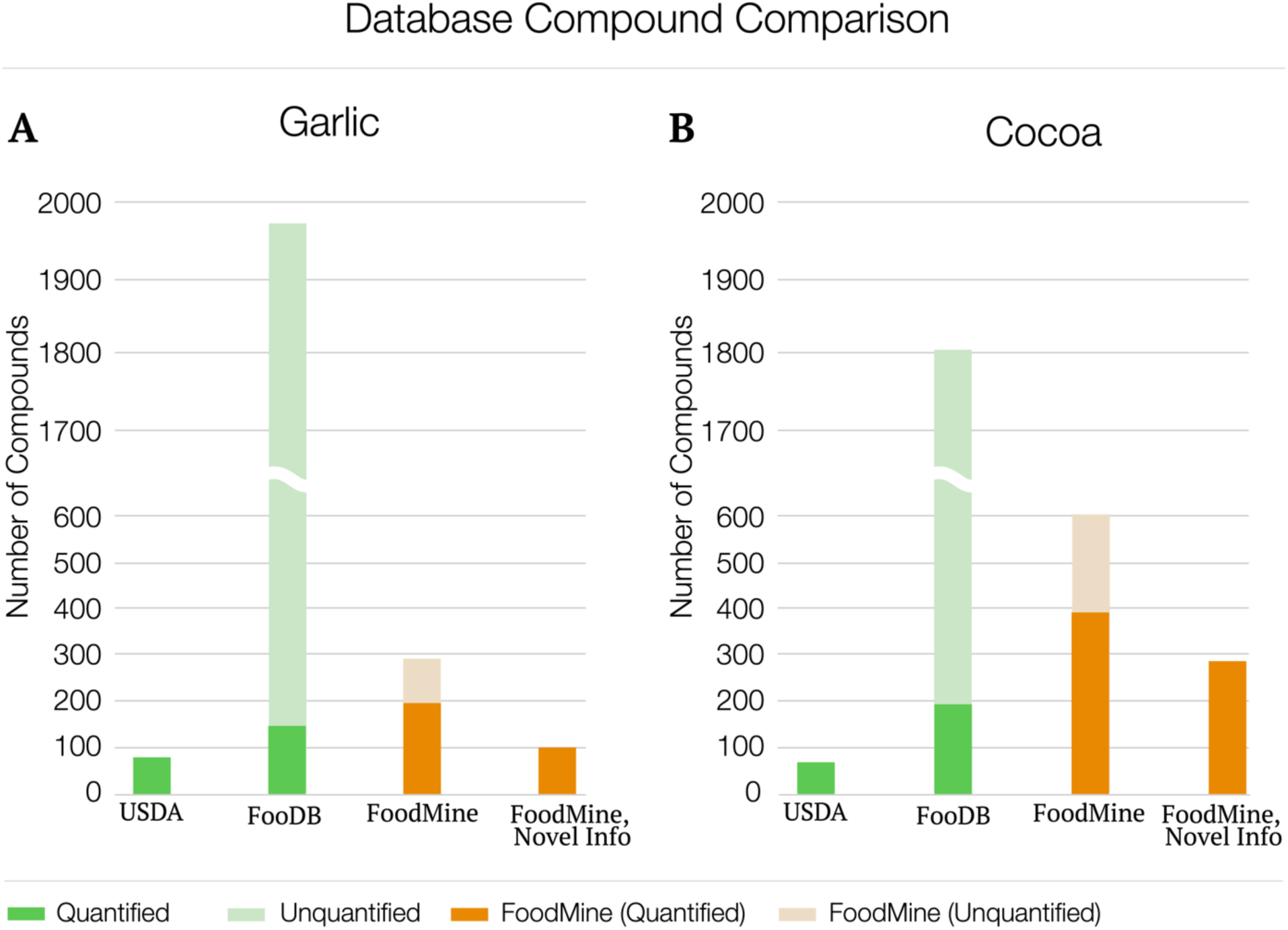
Number of Unique Compounds Recovered by FoodMine, USDA, and FooDB. The plots show the number of unique compounds reported by USDA, FooDB, and FoodMine. The columns display 1) the total number of unique quantified compounds in each database, 2) the total number of unique unquantified compounds in each database, and 3) the number of quantified compounds retrieved by FoodMine and never reported before in USDA or FooDB.

The most frequently reported compounds (Figure 3) in FoodMine are known to play important roles in health effects and flavor. For example, diallyl disulfide is known to contribute to garlic’s smell and taste. More importantly, it is implicated in the health effects of garlic, in particular garlic allergy.^23,24^ Yet, neither USDA nor FooDB offers quantified information for the compound. This is not an isolated case, as Figure 3 shows FooDB and USDA lack information on other frequently explored compounds as well. The need to systemically characterize the nutrient profile of a large number of food items, as USDA does, misses information on those compounds that are specific to a few individual foods, despite the potential role they play in health. Indeed, three of the top ten compounds for cocoa are not quantified in FooDB and one is not listed, while for garlic, five of the top ten compounds are not quantified.

**Figure 3:**
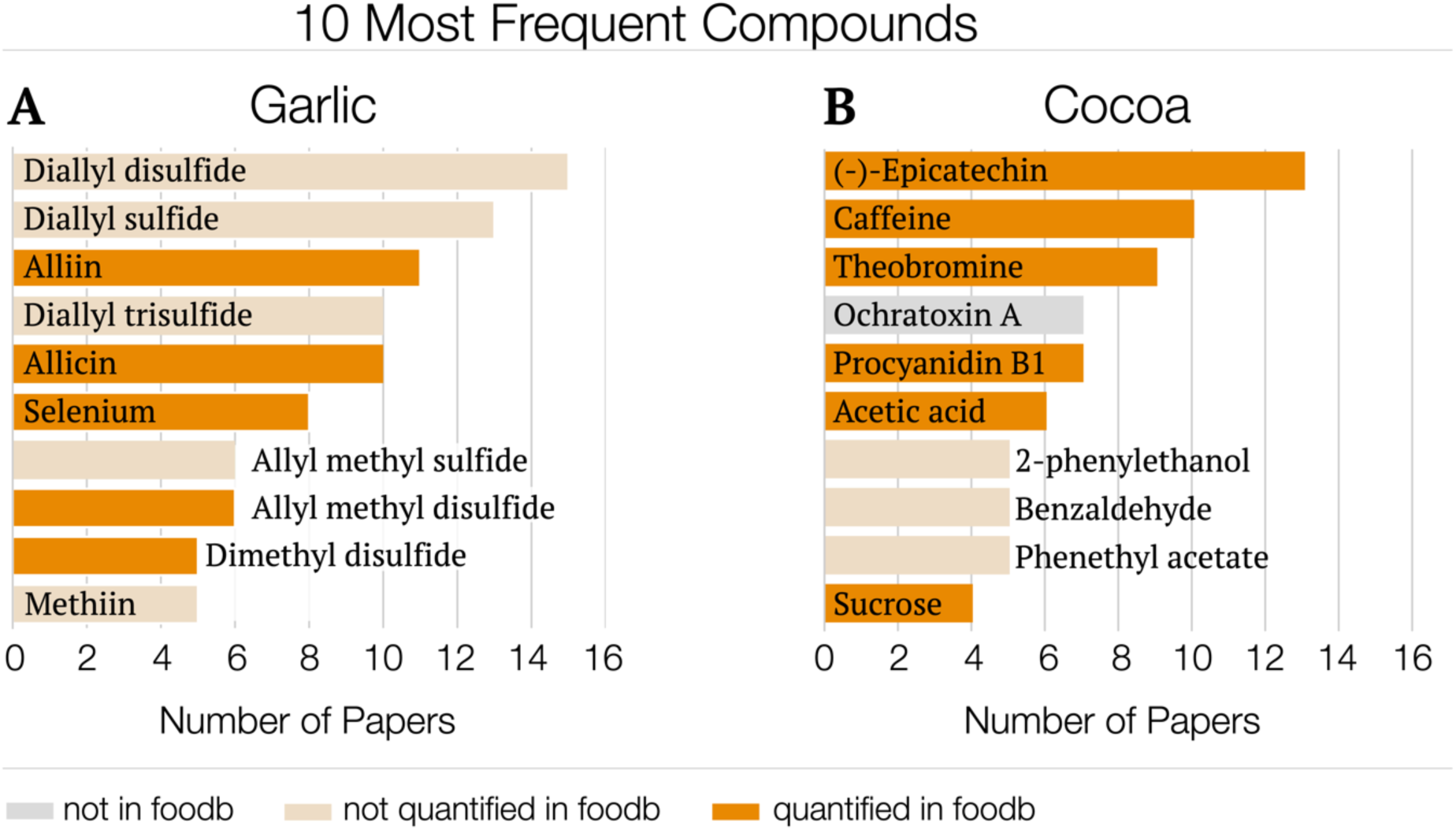
Most Frequently Occurring Compounds in FoodMine. The graphs show the top 10 most frequently occurring compounds in terms of number of recovered papers for (A) garlic and (B) cocoa, gauging the research interest in each product. The y-axis displays the compound name, and the x-axis shows the number of papers that contain records for the given compound.

To understand the accuracy of the collected quantified data, we compared the FoodMine compound measurements to their corresponding values in USDA, the gold standard for measurement reliability among national food composition databases. Given the limited nutrient panel reported by USDA, we were able to compare only 11% of the chemical compounds we recovered for garlic, and 5% for cocoa. The recovered information spanned a full spectrum of molecules, mixing compounds with both small and large relative quantities (Figure 4). Overall, we find a good agreement between the FoodMine-recovered and the USDA-reported values (see Supplementary Material Section 3 for statistics). Garlic has a logarithmic R-squared value of 0.82, indicating a notable correlation between the known amounts and the FoodMine records, while cocoa only reached 0.56. The lower correlation for cocoa is due to a group of amino acids, reported by papers that examined the contents of roasted cocoa, a processing step that alters the quantities of many chemicals, potentially explaining the difference from the USDA measurements.^18,19^ If we remove the data pertaining to roasted cocoa, the logarithmic R-squared increases to 0.75.

**Figure 4:**
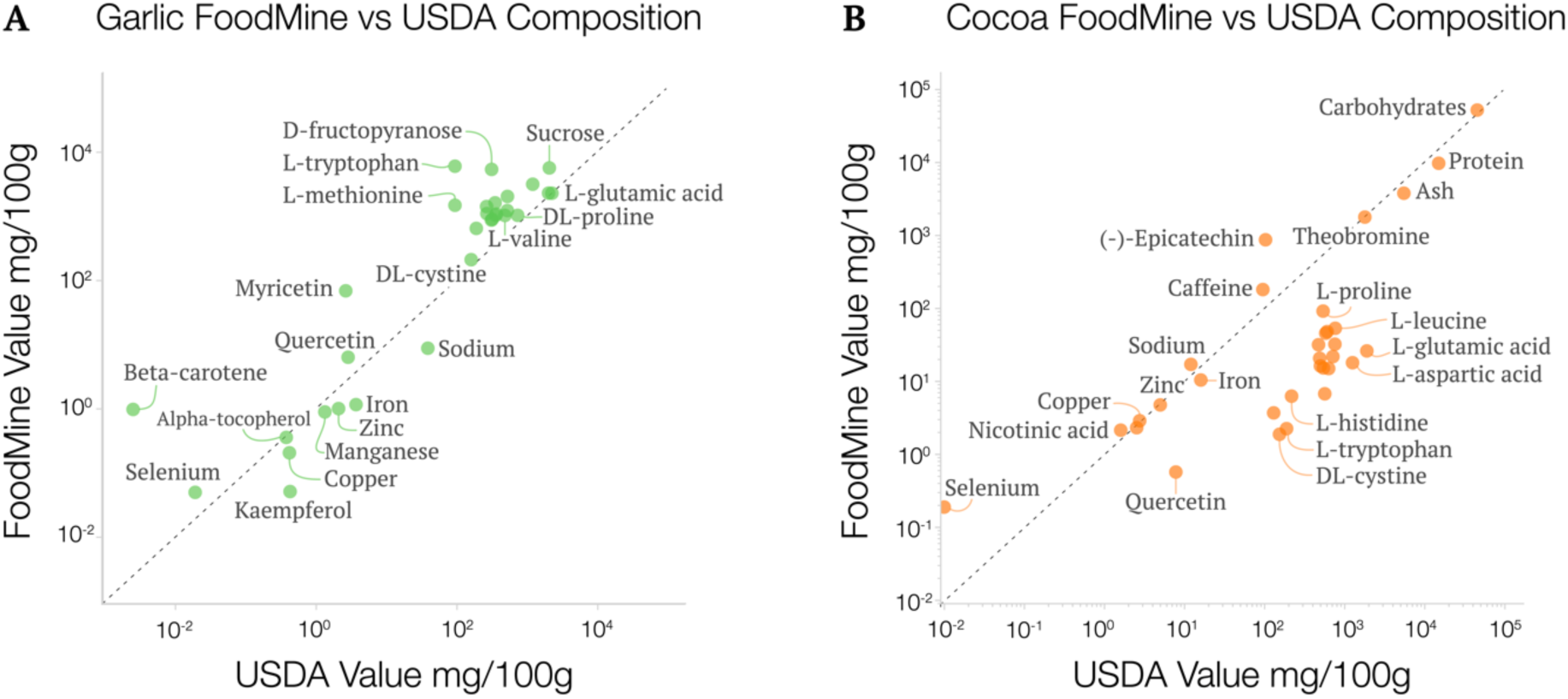
Measurement Comparisons Between FoodMine and USDA. The nutrient concentrations reported by USDA (x-axis), plotted against the content values of matching compounds in FoodMine (y-axis). The dotted line represents the diagonal. We excluded three and two compounds for (A) garlic and (B) cocoa, respectively, because USDA reported zero values for those compounds.

To offer a more comprehensive understanding of the classes of chemicals we retrieved and their relation across different databases, we created chemical embeddings using the unsupervised machine learning tool Mol2Vec.^25^ Chemical embeddings capture the structural similarity of the chemicals in a low dimensional space. Indeed, as shown in Figure S5, when the chemical classification is known, chemicals belonging to the same class tend to be closer in the embedding space defined by Mol2Vec, suggesting that chemical embeddings successfully capture structural information. This process maps the structural knowledge of the compounds we retrieved, and can be used to integrate further information characterizing the compounds. For instance, given the interest in the association between food-borne chemicals and health outcomes, we can leverage the Comparative Toxicogenomics Database (CTD), that reports manually curated associations between chemicals and diseases.^26^ After matching the total number of health implications to each of the quantified chemicals in FoodMine, FooDB, and USDA, we layered this information on the obtained Mol2Vec embedding (Figure 5). We find that FoodMine covers more chemicals with health effects than FooDB and USDA (see Figure 5A vs B and C, D vs E and F), a difference particularly clear for cocoa (D vs E and F). Furthermore, we find that the chemicals with health associations are more evenly dispersed throughout the embedding space for FoodMine, implying that FoodMine captures chemicals from rather different chemical classes. Overall, the FoodMine cocoa pilot has detected noticeably more organic, benzenoid, and hydrocarbon compounds, as seen by the absent spaces in E (USDA) and F (FooDB) compared to D (FoodMine) (see Supplementary Material Section 3). In summary, compared to the existing databases, FoodMine detects more chemicals with health associations, distributed over a wider range of chemical classes, reflecting a selection bias in the literature: the research community appears to be more focused on chemicals with known health outcomes. Interestingly, there is no overlap between the papers contributing to FoodMine and those manually curated in CTD, meaning that we are recovering information from multiple scientific communities, not only health sciences (see Supplementary Material Section 3).

**Figure 5:**
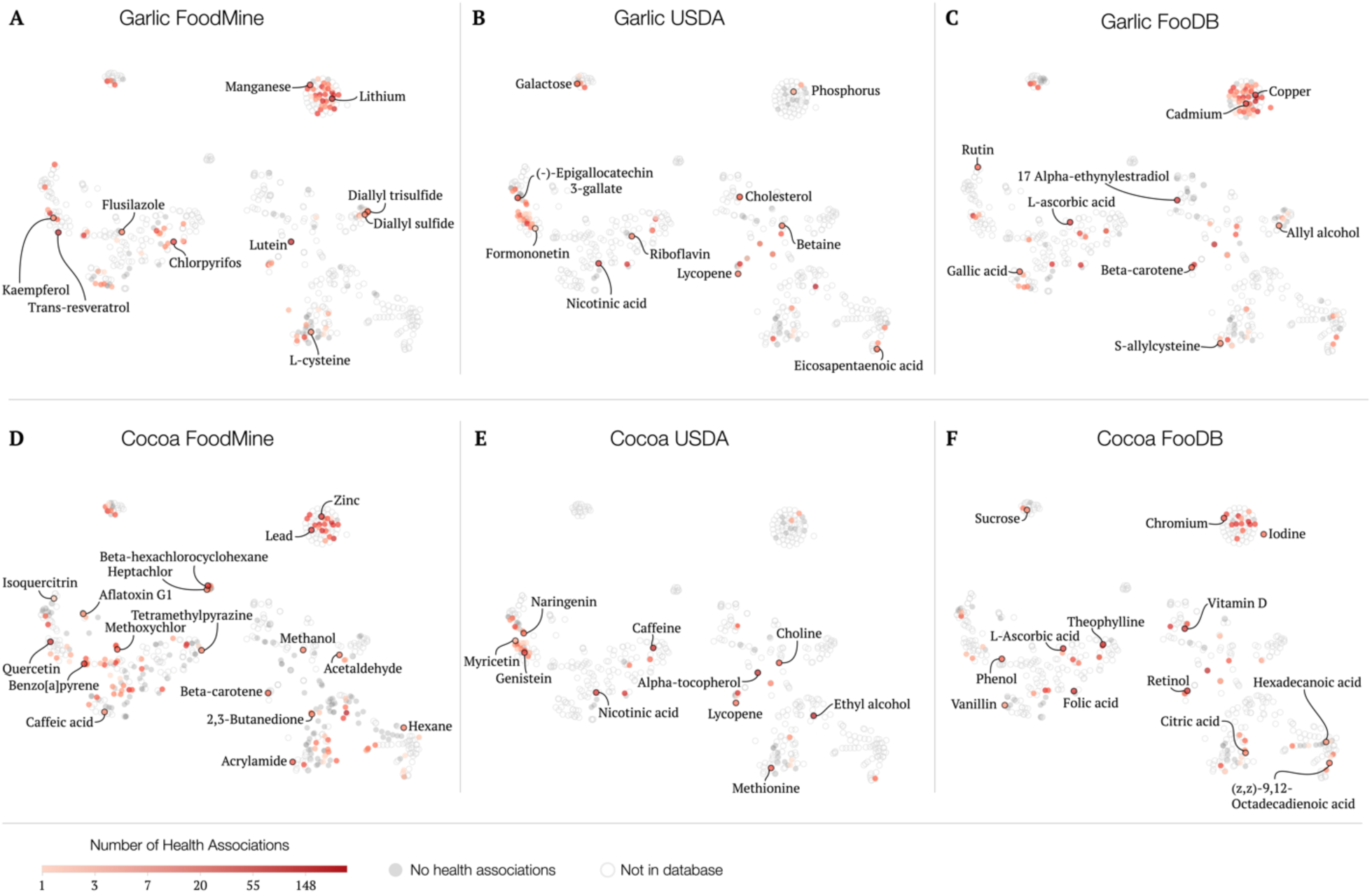
TSNE Dimensionality Reduction of Chemical Embeddings with Health Associations. TSNE plots of Mol2Vec chemical embeddings for garlic (A, B, and C) and cocoa (D, E, and F). The colors of each data point encode the number of health implications associated with the compounds based on the CTD database. Dark gray represents chemicals with 0 health associations. We show chemicals catalogued by each studied database for FoodMine (A & D), USDA (B & E), and FooDB (C & F). Markers are filled if the database contains the chemical, and empty if it does not.

## Discussion

Our knowledge pertaining to the more than 26,000 chemicals expected to be present in food, as reported in various databases, is highly incomplete. This incompletion inspired our efforts to examine how much additional uncatalogued knowledge is scattered in the scientific literature. The invisibility of these compounds to experimental, clinical, epidemiological, and demographic studies – the virtual “dark matter” of nutrients – represents a major roadblock towards a systematic understanding of how diet affects our health. The introduced FoodMine pilot systematically scanned the scientific literature, identifying information about a large number of novel, quantified compounds reported by individual papers. We find that the collected information considerably extends our understanding of food composition. Furthermore, many of the recovered compounds have direct relevance to health and nutrition. For instance, the sulfides, quantified by FoodMine, are responsible for garlic’s unique health effects, yet are currently not quantified in USDA or FooDB.

Garlic and cocoa are only two of the over a thousand natural foods commonly consumed by humans, hence our study supports the hypothesis that there is abundant information in the literature on the composition of other ingredients as well. Indeed, the search terms we used in FoodMine to retrieve papers from PubMed were narrow, and the selection of papers we manually evaluated is small compared to the total body of potential knowledge present in the literature. Consequently, there is likely additional information for garlic and cocoa, not yet captured by FoodMine. Other search terms, focusing on compound classes rather than foods, could uncover an additional body of information about the chemical composition of these ingredients, knowledge that can be generalized to other ingredients as well. For instance, by targeting ‘NEPP’, i.e. non-extractable polyphenols, FoodMine could in principle collect and disambiguate the available literature that reports the food content of this class of chemical compounds, often overlooked by food databases, despite the growing interest on their interaction with the human gut microbiome.^27–29^

Our efforts for garlic and cocoa have proven the existence of a sizable yet scattered literature pertaining to their chemical composition, offering a consistent gain of compositional information compared to what is currently available in food databases. With our pilot we focused on chemical information that has been measured by scientists but was effectively lost to the public, due to the lack of storage and disambiguation in accessible databases. Indeed, despite the complexity characterizing the nutrient dark matter, the food consumption is still far from the efforts of genomic and proteomic research in the construction of biobanks and consortiums, curating and storing the chemical compounds identified in food. Documenting what is currently known about food composition is a necessary step towards further experimental efforts. In this perspective, the output of FoodMine constitutes a valuable starting point for the creation of standards necessary for targeted metabolomics, helping identify and quantify the variability of these chemical compounds in food.^30,31^

Our next goal is to expand the data collection to multiple staple ingredients. We are prioritizing our search according to the consumption and production statistics available in national and international surveys like NHANES^32^ and FAOSTAT^33^, aiming to target foods that would help drastically improve the chemical coverage of our diet, and benefit health studies. While manual curation is still needed to extract measurement details from papers, our machine learning algorithm is ranking the papers in order of relevance, to expedite the data collection. Given the heterogeneous scientific language used to describe food, the second phase of this pilot is key for acquiring additional data training to learn novel language features, such as the occurrence of particular n-grams^34,35^, to maximize the applicability of the algorithm to different foods, without losing precision.

## Methods

Literature mining consisted of three steps: search, selection, and information extraction. We began by searching PubMed with the search term ‘garlic’ and ‘cocoa’, using the Pubmed Entrez Programming Utilities API.^36^ After retrieving the PubMed ID’s for search results, we again used the API to retrieve information for each PubMed entry associated with the PubMed ID. We used text matching to scan each PubMed entry’s MeSH terms and abstract for words relevant to biochemicals, food, and pre-selected measurement methodologies, after the API query (see Supplementary Material Section 1). The algorithm filtered 5,676 results for garlic and 7,620 papers for cocoa to 415 and 475 results, respectively. We collected papers from the “Full text links” of the combined 900 entries. We skipped an entry if it did not list any “Full text links” or we did not have access to the paper associated with the links, recovering papers for 299 of the 415 PubMed entries for garlic, and 324 of the 475 entries for cocoa. We also spot-checked search results that fell outside the assessment criteria to quantify the effectiveness of the filtration step. Of those, 0/10 papers contained relevant information for garlic and 3/10 papers contained relevant information for cocoa, indicating that the filtering did eliminate some papers with potentially relevant information.

Papers were individually read by a human assessor to determine whether or not they contained information on the chemical composition of garlic or cocoa. The assessment fell into three categories: “not useful” (papers not containing relevant information for chemical contents in food), “quantified” (papers containing quantitative chemical composition that could be translated to unveil its precise contents in samples), and “unquantified” (paper containing chemical composition, but it could not be converted to unveil precise quantities). Examples of unquantified results were compounds detected by a mass spectrometer that only reported relative percentages, hence we could not record their contents in garlic or cocoa. In addition to the human mining procedure, we used machine learning to create a paper classification algorithm, helping to automatize future data collection (See Supplementary Material Section 1). This algorithm takes as input the filtered samples and predicts which papers will contain information on the chemical content of cocoa or garlic. We applied SMOTE sampling technique to balance the labeled data, as only 27% of the reviewed papers contained information on the chemical content of cocoa or garlic.^37^ Our algorithm achieved an f1 score of 75.5% on the testing set, better than random. These results are in spite of a limited training set, and could improve with more labels from other foods beyond garlic and cocoa.

All records for a single unique compound were merged into a single entry by calculating the mean of quantified record values. As different papers use different variations of a compound’s name, we applied a chemical disambiguation scheme using PubChem CIDs to add keys to the compounds (see Supplementary Material Section 2).^38^ For each entry, we reported the average content value across all data points standardized in units of mg/100g, and captured additional statistics, such as the highest and lowest reported measurement of the chemical, variance across measurements, and number of measurements. Finally, we leveraged the PubChem CIDs to retrieve a string representation of the structural properties of the molecule (chemical SMILE) which we used as the input for Mol2Vec. Once we learned the vector representation for each chemical, we further reduced the dimensionality using TSNE to obtain the maps shown in Figure 5 and Figure S5.^39^

## Data Availability

The raw data and processing code are available at https://github.com/fhooton/FoodMine.

## Additional Information

See Supplementary Material for more information.

## Acknowledgements

We thank Daniela Barbery-Flambury for the manual data collection and Dr. Michael Sebek for advice on compound classes.

## Author Contributions

FH performed data analysis, programming, and contributed to writing the manuscript. GM designed the project procedure and analysis, and contributed to writing the manuscript. ALB contributed to interpreting the results and writing the manuscript. GM and ALB conceived the project. FH and GM contributed equally to the project.

