## Supplementary Material for "Exploring Food Contents in Scientific Literature with FoodMine"

#### Section 1: Data Collection

Literature mining consisted of three technical steps: search, selection, and information extraction. We began with a general search query, and then applied programmatic filters to the resulting query. After each filter, we inspected the remaining papers to determine if they contained any relevant information. If the paper was selected, we manually extracted information pertaining to compounds reported for garlic and cocoa, and organized it in a central database. We chose PubMed as the search engine for this process, as it represents the largest biomedical database that contains information for over 28 million research papers, and uses open paper indexing with professional review through MeSH terms.

Given the recent interest in milk composition determined by experimental research combined with literature investigation, we explored PolySearch2, an online tool to search free-text corpora such as PubMed Central.[1,2] We used PolySearch to retrieve information relevant to garlic and cocoa by entering "garlic" and "cocoa" as search terms, respectively. We searched for relevant food metabolites, MeSH compounds, and toxins. PolySearch2 generated 50 results across the three categories for garlic, and 24 results for cocoa. While PolySearch found chemicals associated with garlic and cocoa, our FoodMine process indicates that more information is available. Furthermore, PolySearch did not retrieve compositional information for the displayed chemicals, an impediment to its direct application in health and nutrition.

#### Querying PubMed

We used the search function from the PubMed Entrez Programming Utilities API to search the database, and extracted the PubMed IDs of all entries returned from the search query in an XML format.[3] Once we had a list of search IDs, we again leveraged the PubMed API to retrieve the title, year, abstract, MeSH Terms, and Qualifier Terms for each document using the python package *lxml*.

URL to search PubMed: https://eutils.ncbi.nlm.nih.gov/entrez/eutils/esearch.fcgi?db=pubmed&term= + search terms

URL to retrieve info by PubMed ID:

https://eutils.ncbi.nlm.nih.gov/entrez/eutils/efetch.fcgi?db=pubmed&id= + PMID

Example Entry:

https://www.ncbi.nlm.nih.gov/pubmed/?term=Molecular+definition+of+the+taste+of+roasted+cocoa+nibs+(Theobroma+cacao)+by+means+of+quantitative+studies+and+sensory+experiments

Example XML Information:

https://eutils.ncbi.nlm.nih.gov/entrez/eutils/efetch.fcgi?db=pubmed&id=16848542&retmode=xml

See pubmed_util.py for code, at https://github.com/fhooton/FoodMine.

#### Preliminary Entry Filtration

Each PubMed ID, associated with a specific paper, brings information on the title, authors, year, abstracts, and MeSH terms. We filtered search results by checking if the abstract or MeSH terms contained at least one word from a ‘food’, ‘chemical’, ‘scientific name’, or ‘general’ dictionary, and that the abstract or MeSH terms contained at least one of the following measurement methods: 'spectrometry', 'chromatography', or 'spectrophotometry'. We did not manually review entries not meeting these two conditions.

We created the ‘food’, ‘chemical’, and ‘scientific name’ dictionaries from information listed on the FooDB website. Specifically, we scraped all distinct food names, chemical names, and scientific names and stored each as a separate list. The ‘general’ dictionary was designed for papers with more general descriptions that might contain valuable information, and lists words such as ‘food’, ‘meat’, ‘vegetable’, and ‘database’.

#### Summary of the Automatic Data Collection Process

We summarize here the key steps represented in Figure 1, Box `Automatic’:

1. Retrieve abstracts and metadata from PubMed using the search terms “garlic” and “cocoa”, independently.
2. Filter out papers that do not contain any terms in the food term dictionaries or target measurement methods.

#### Manual Review

After the automatic filtering, we undertook an extensive manual identification and extraction procedure. First, we identified if a paper had relevant information. Some experiments modified the contents of chemicals to see their effect on food. In that case, we did not consider the modified chemical contents, but only the control values for them. Cooking methods that did not artificially inject specific compounds, like baking, were included. The set of criteria we developed to standardize the data collection and classify the measurements as pertinent is shown in Table S1. We read the entire research paper for information on food contents. Most food content information came from tables or graphs, but we also found information in the paper text. If we did find relevant content information, we checked the surrounding text and the methodologies section to verify the experiment did not alter the foods. Experiments often spiked foods with various chemicals to observe the change in contents over time. We also determined if compounds were “quantified” or “unquantified”. Quantified compounds could be converted into an absolute mass, like mg/100g. Unquantified compounds had relative mass measurements, or only recorded a chemical's presence. Studies using solely mass spectrometry to measure relative chemical quantities produced many unquantified compounds.

We aimed to gather several types of information from research papers. Table S1 describes the target information we extracted. We used several tools to speed up the extraction process for quantitative data. For instance, if a paper presented quantitative data in a table, we used Tabula to automatically extract the table.[4] PDF's with tables as images were not always detected, and we extracted the data manually in those instances. For graphs, we used Automeris to extract an approximation of the value. We extracted all information into CSV files.[5]

When we extracted information in the quantified category, we recorded all units of mass the paper reported, such as micrograms of a compound per 100 grams. The quantified category also contained units that could be calculated to find the absolute mass, such as micrograms per liter of solution. We calculated these values using the mass of garlic or cocoa added to the solvent in the experimental method of the corresponding papers. If a paper listed a value with a unit involving molarity, like micromoles per liter, we used the molecular weight of each compound to calculate the mass. We also recorded content values of "0" and "not detected" for compound contents tested but not detected in garlic and cocoa. We stored the extracted information as records in a csv format, and left cells blank if a paper did not provide information for the qualitative categories. Each record is a specific data point containing the information categories, and papers could have multiple records per compound if the methodology prepared samples differently, or listed independent sample batches.

#### Modeling

To accelerate relevant paper curation, we leveraged the manual efforts of our analysis to curate training data for machine learning approaches. In this study, we experimented with a model that would label papers containing information on the chemical composition of food. In our experiment, we explored logistic regression, support vector machine, random forest, XGBoost, decision tree, neural network, and k nearest neighbors models. Our first experiment yielded poor results due to class imbalance, as 73% of the labeled papers do not contain information pertaining to chemical composition. We then applied the synthetic minority over-sampling technique (SMOTE) to balance the classes,[6] and with XGBoost we achieved an average f1-score of .755 (.74 for papers that did not contain chemical compositional information and .77 for those that did). Using training on two foods to decrease overfitting, and a third food as the holdout set, could improve the model in the future.

See filter.py for code, at https://github.com/fhooton/FoodMine.

#### Section 2: Chemical Disambiguation

Many compounds did not perfectly match between FoodMine, USDA and FooDB (sql data dump dated 06/29/2017). To compare the different chemical descriptions, we needed to disambiguate the chemical nomenclature, leveraging the PubChem database to address the naming differences, a large database of chemical compounds from NCBI that contains information on compound synonyms, grouped under unique identifiers, called CIDs.[7] For this purpose, we queried all compound strings for FoodMine, FooDB, and USDA to retrieve the associated CIDs and inChI keys. By selecting the first 14 characters of the inChI key, we determined the code for the structure of each chemical compound. Chemicals with identical structural codes were considered identical and removed, thereby filtering to unique chemical structures and omitting information regarding stereochemistry and ionization. By restricting the assessment to the physical structure, the varying methodologies between the databases with respect to stereochemistry and ionization were removed, allowing for objective comparisons between databases. If no inChI key was found, we proceeded by string matching. After these steps, if a FoodMine compound did not match any FooDB and USDA compounds, we concluded that it was unique to FoodMine.

PubChem Example: https://pubchem.ncbi.nlm.nih.gov/compound/65036

See Data_Statistics.ipynb and collected_data_handling.py for code, at

https://github.com/fhooton/FoodMine.

### Section 3: Data Analysis

We assessed the validity of our data collection with summary statistics, comparison with state-of-the-art food composition databases, and chemical embedding visualization. When comparing FoodMine to USDA and FooDB, we used multiple categories for each source to synthesize a more representative sample. The USDA categories used are “Garlic, raw” and “Spices, garlic powder” for garlic, and “Oil, cocoa butter”, “Cocoa, dry powder, Hershey's European style cocoa”, “Cocoa, dry powder, unsweetened”, “Cocoa, dry powder, unsweetend, processed with alkali”, and “Cocoa, dry powder, hi-fat or breakfast, processed with alkali” for cocoa. The FooDB categories used are “Garlic” and “Soft-necked Garlic” for garlic, and “Cocoa Bean”, “Cocoa Butter”, “Cocoa Powder”, “Cocoa Liquor” for cocoa.

#### Basic Statistics

After disambiguating the compounds from FoodMine, we computed some summary statistics. Figure S1 shows the distribution of total chemical records per paper for (A) garlic and (B) cocoa, grouped by papers containing quantified and unquantified chemical information. We also included the distribution of unique, or distinct, chemicals per paper in (C) garlic and (D) cocoa, again grouped by papers containing quantified and unquantified information.

#### Database Comparison

In addition to Figure 2, Table S2 and S3 show the numerical comparison between FoodMine and the aggregate of FooDB and USDA. We have listed the calculations for the percentage of quantified chemicals present in FoodMine and the aggregate of FooDB and USDA, and the percentage increase in quantified information.

Percentage of Quantified (USDA U FooDB) Chemicals

- Garlic: 177 quantified chemicals / 1,920 total chemicals ≈ 9.22%

- Cocoa: 128 quantified chemicals / 1,853 total chemicals ≈ 6.91%

Percentage of Quantified FoodMine Chemicals

- Garlic: 201 quantified chemicals / 289 total chemicals ≈ 69.55%
- Cocoa: 395 quantified chemicals / 598 total chemicals ≈ 66.05%

Percentage of Novel Quantified FoodMine Chemicals

- Garlic: 96 / 201 quantified chemicals ≈ 47.76%
- Cocoa: 283 / 395 quantified chemicals ≈ 71.65%

Percentage Increase of Quantified Information

- Increase for garlic: 96 novel quantified FoodMine chemicals / 177 quantified (USDA U FooDB) chemicals ≈ 54.24%
- Increase for cocoa: 283 novel quantified FoodMine chemicals / 128 quantified (USDA U FooDB) chemicals ≈ 221.09%
- Average Increase in information = (54.24% + 221.09%) / 2 ≈ 137.66%

#### Sample Characteristics

As researchers investigate food for a variety of purposes, samples in research literature show different features such as processing methodology, part of food, or physical state. Garlic and cocoa, widely used in different recipes, can be reported as powder, oil, bean, or otherwise. Figure S3 shows the percentage of papers containing the corresponding sample characteristic. Among the features listed, clove most frequently occurs for garlic and powder for cocoa. These characteristics are not mutually exclusive, as a sample description could be “fresh garlic clove” or “raw cocoa powder”. Unfortunately, some papers do not list the features characterizing the samples. The inconsistent characterizations of garlic and cocoa samples highlights a gap in research communications, and limits our ability to do granular comparisons across different sample collections.

#### Comparison with Phenol Explorer

Phenol Explorer is a database containing food polyphenol contents extracted from scientific literature, and represents an earlier effort in food literature mining. The database curated academic papers on polyphenols using combinations of specific food and polyphenol search terms over the closed-source FSTA database (Food Science and Technology Abstracts), and manually evaluated each result.[8,9] For instance, as reported in [8], the query for carrot polyphenols followed as: "[(carrot or carrots) and (vitisin or resveratrol or stilbene* or polyphenol* or phenol* or flavon* or flavan* or cinnamic* or benzoic* or anthocyan* or quercetin or luteolin or myricetin or apigenin or isorhamnetin or catechin* or epicatechin* or epigallocatechin* or nar* or hes* or lignan* or tannin* or ellagitannin* or ellagic or kaempferol or proanthocyan* or procyan* or caffeic or ferulic or sinapic or chlorogenic) and (composition* or compound or compounds or content or contents or determin* or quanti* or profil* or identif*)]". Figure S4 shows the comparison between the number of unique compounds listed in Phenol Explorer and FoodMine. As Phenol Explorer contains 5 compounds for garlic and 35 for cocoa, the number of compounds in FoodMine well exceeds Phenol Explorer for both food pilots. FoodMine recovered 14 of the 35 compounds in Phenol Explorer for cocoa, and none of the compounds in Phenol Explorer for garlic. The chemicals reported by Phenol Explorer were collected from 7 papers for garlic (4 available on PubMed), and 8 papers for cocoa (7 available on PubMed). Interestingly, FoodMine recovered part of these compounds from a completely orthogonal literature, suggesting that our results could be further extended by adding specific search terms for compounds of interest, as mentioned in the Discussion Section.

#### FoodMine-USDA Fit Statistics

For both garlic and cocoa we fitted the relation $\log y = \alpha + \beta \log x$, to compare the chemical contents retrieved by FoodMine ($x$) with those reported in USDA ($y$). For garlic we find $\beta = 0.84(0.692, 0.991), \alpha = -0.228\,(-1.131, 0.675)$ and $R^2 = 0.821$, showing a weak sublinearity. For cocoa, after removing the outliers for roasted cocoa, we find $\beta = 0.92(0.584, 1.263), \alpha = 1.103(-0.455, 2.662)$ and $R^2 = 0.745$. These results are consistent with the identity relation $y = x$, however, the conclusions are limited by the small overlap between FoodMine and USDA.

#### Molecule Embedding and TSNE

The molecular embedding technique used in the paper originated from Mol2Vec.[10] We retrieved chemical SMILEs for all the quantified chemicals across USDA, FooDB, and FoodMine from PubChem. Because a compound needed to have a PubChem CID to be queried for the chemical SMILE, this analysis only includes compounds with PubChem CIDs. The compounds from all databases were combined into a single list of unique compounds. Once we retrieved SMILEs for the whole compound list, we used the RDKIT python package to calculate Morgan Fingerprints of radius 1.[11] We transformed the substructure code of the fingerprint into a string of substructure codes, ordered according to the preset order of the fingerprint. The substructure codes represented words and the fingerprint strings represented sentences for Mol2Vec. After forming the fingerprint strings, we replaced a substructure code with 'UNK' if it appeared 3 times or less throughout all chemicals. We used the Word2Vec class from the gensim python package, training a skip-gram word embedding model with embedding size of 100.[12] We created a fingerprint vector per compound by averaging the embeddings for all substructure vectors. Finally, we used the TSNE function from the sklearn python package to reduce the fingerprint vectors to two TSNE components.[13]

To characterize the type of compounds in our project, and assess if the TSNE embedding captured relevant structural information, we matched the compounds with their respective FooDB chemical classes. Figure S3 shows the TSNE embedding for compounds in FoodMine, FooDB, and USDA. The plots verify that meaningful information is captured, as chemicals belonging to the same class tend to be closer in space, suggesting that the chemical embeddings are successfully capturing structural information.

We used the number of manually accrued health associations from CTD to compare health consequences across FoodMine, FooDB, and USDA. For this analysis, we simply counted the number of "markers/mechanisms" or "therapeutic" labels per chemical. However, we wanted to ensure that there was no selection bias towards FoodMine papers. We automatically queried all CTD chemicals in PubChem and attached PubChem CIDs, and selected paper PMIDs for the compounds that matched FoodMine chemical CIDs. There was no overlap between these CTD PMIDs and the PMIDs from FoodMine. As an extended step, we also used the Microsoft Academic Graph to retrieve the citations for FoodMine papers, and queried each citation title in PubMed to retrieve the respective PubMed PMIDs.[14,15] Only two citation PMIDs overlapped with the CTD PMIDs.

See Molecule_Embedding.ipynb for code, at https://github.com/fhooton/FoodMine.

### Figures and Tables

| Category | Description |
|---|---|
| Food Item | Food item to be included in the database (for example 'Garlic' for this study) |
| Part | Part of the food a paper analyzes (for example bulb, root) |
| Physical State | Physical state of the food in a study (for example powder, raw plant) |
| Instrumentation | Instrument used in research study (for example Gas Chrom., Mass Spec.) |
| Preparation | Specific preparations of the study's foods (fresh, baked, boiled) |
| Number of Samples | Number of replicates used for samples |
| Compound | Compounds examined |
| Value | Quantified value produced from study |
| Variance | Variance associated with particular value |
| Unit | Units of measure for each repeated value |

***Table S1. Information Categories Used During Data Collection.*** *The qualitative categories listed in the table help evaluate if a paper contained pertinent, useable compound information, and form structured fields for data extracted from papers.*

| | Quantified in FoodMine | Unquantified in FoodMine | Zeros in FoodMine | Not in FoodMine |
|---|---|---|---|---|
| Quantified in (USDA U FooDB) | 68 | 4 | 0 | 105 |
| Unquantified in (USDA U FooDB) | 35 | 6 | 0 | 1,767 |
| Zeros in (USDA U FooDB) | 2 | 1 | 0 | 48 |
| Not in (USDA U FooDB) | 96 | 74 | 3 | |

***Table S2. Comparison of FoodMine with USDA and FooDB with respect to garlic.***

| | Quantified in FoodMine | Unquantified in FoodMine | Zeros in FoodMine | Not in FoodMine |
|---|---|---|---|---|
| Quantified in (USDA U FooDB) | 57 | 11 | 0 | 60 |
| Unquantified in (USDA U FooDB) | 54 | 8 | 0 | 1,634 |
| Zeros in (USDA U FooDB) | 1 | 0 | 0 | 28 |
| Not in (USDA U FooDB) | 283 | 183 | 1 | |

***Table S3. Comparison of FoodMine with USDA and FooDB with respect to cocoa.***

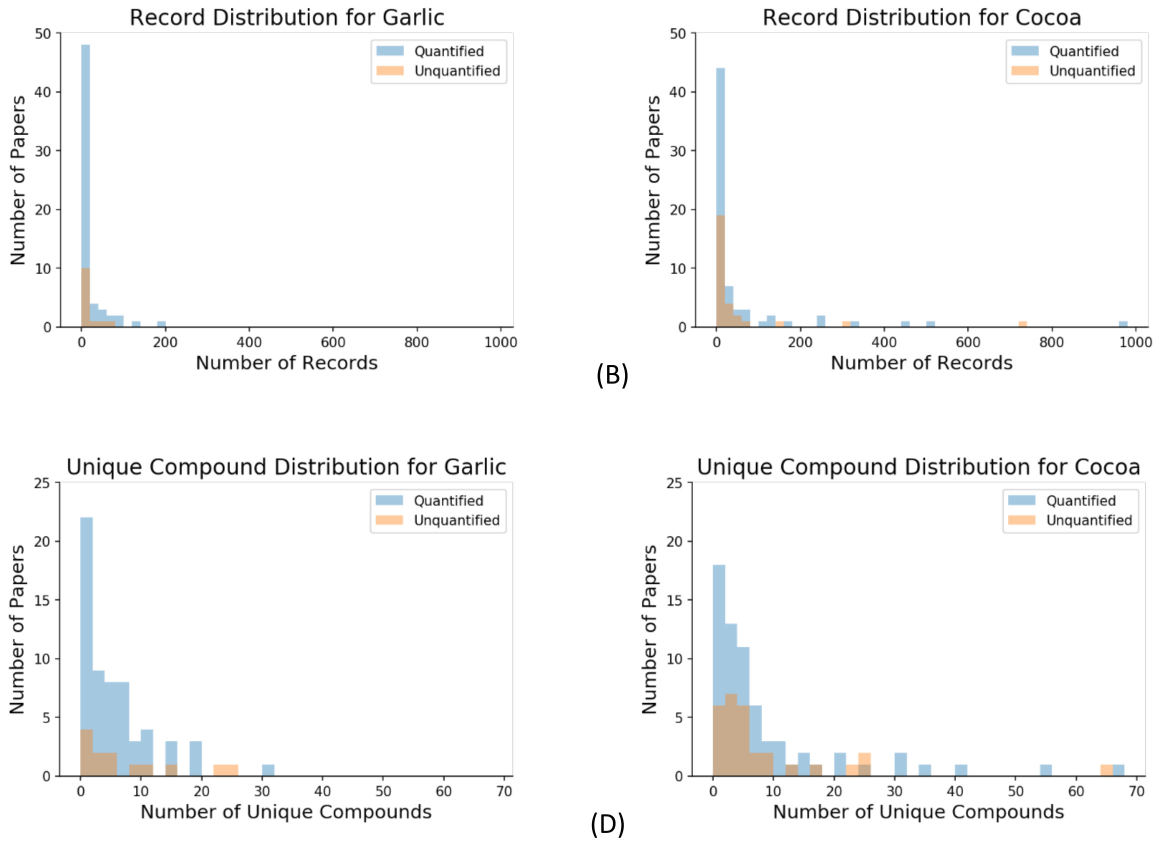


***Figure S1: Record and Unique Compound Distributions in FoodMine.*** *The record distribution (A and B) considers the number of records per paper, while the unique compound distribution (C and D) shows the number of unique compounds per paper.*

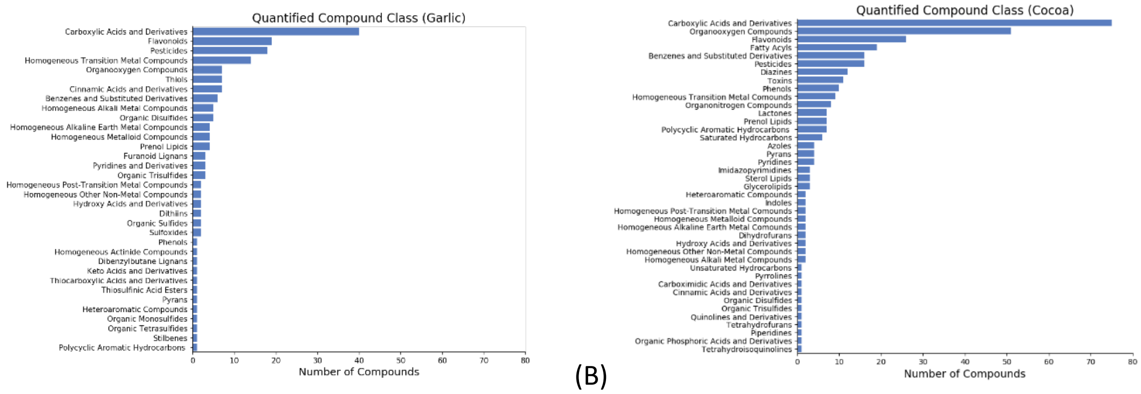


***Figure S2: Number of Compounds in Chemical Classes.*** *The number of unique quantified compounds in each class for (A) garlic and (B) cocoa. We used FooDB to assign chemical classes.*

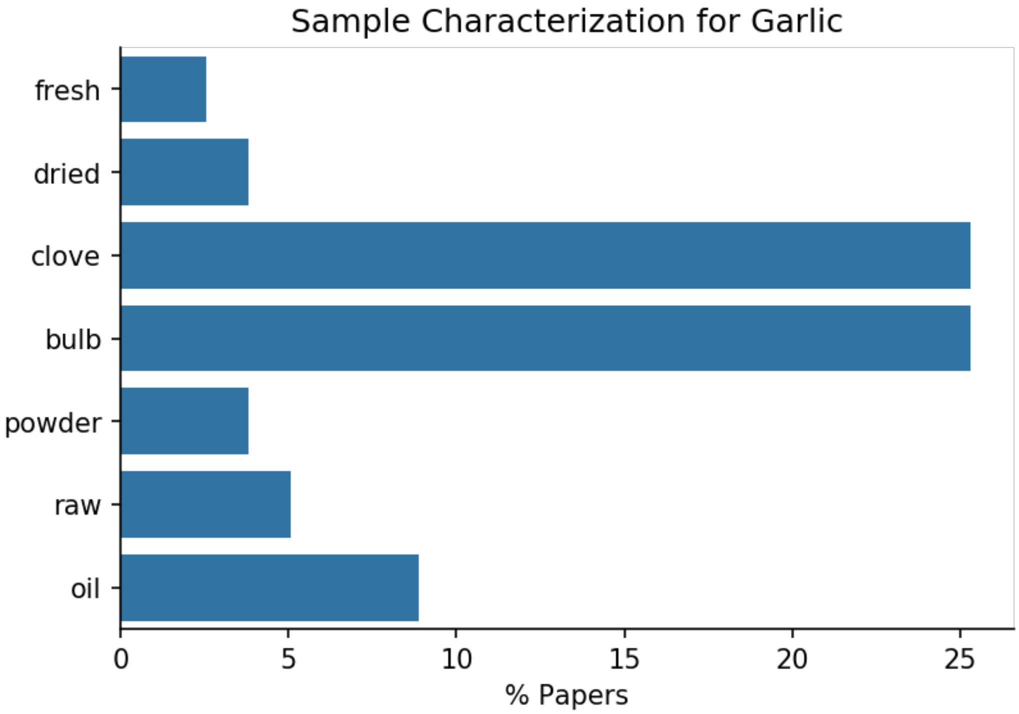


(A)

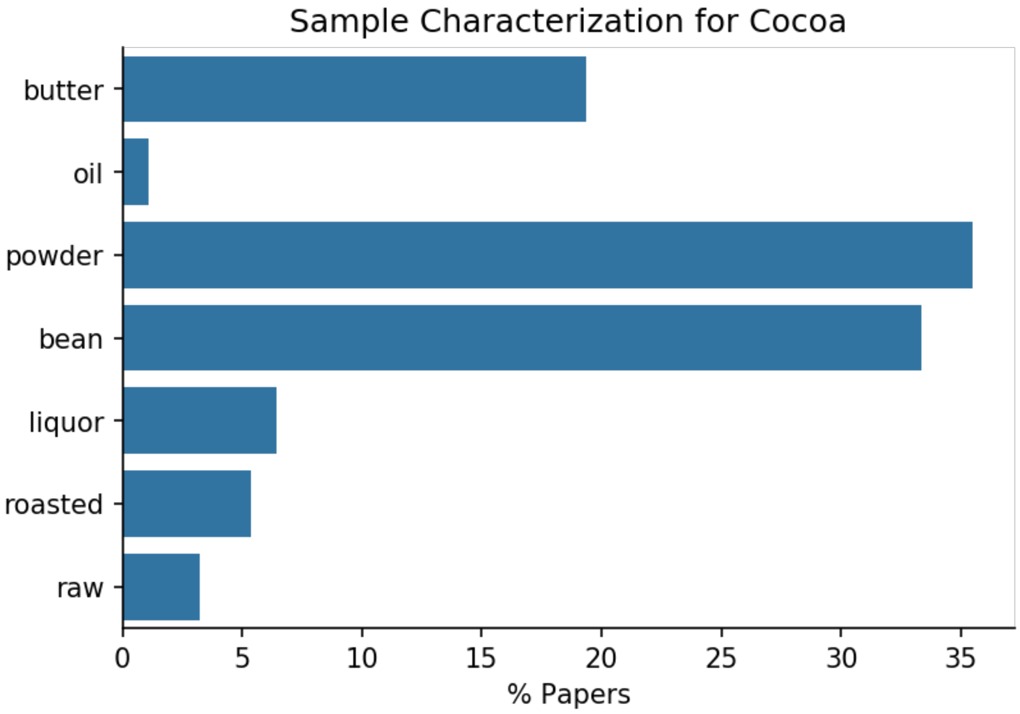


(B)

***Figure S3: Sample Characterization.*** *Percentage of papers with selected features for (A) garlic and (B) cocoa, collected in FoodMine.*

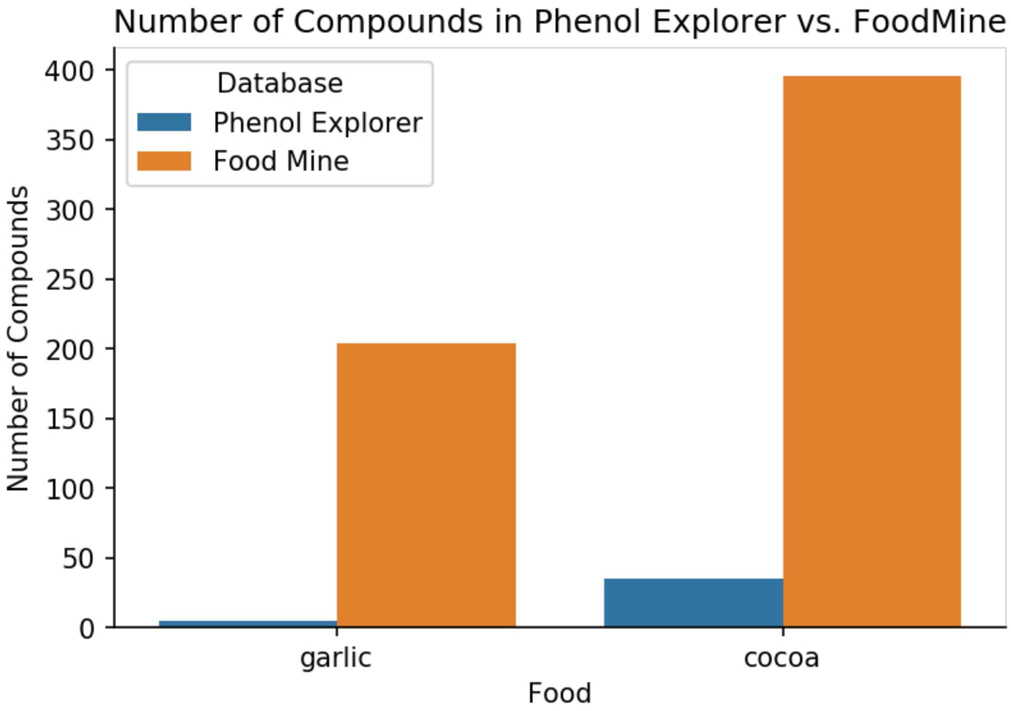


***Figure S4: Number of Unique Compounds collected by Phenol Explorer and FoodMine.*** *Number of unique compound entries listed in Phenol Explorer and FoodMine for garlic and cocoa.*

(A)

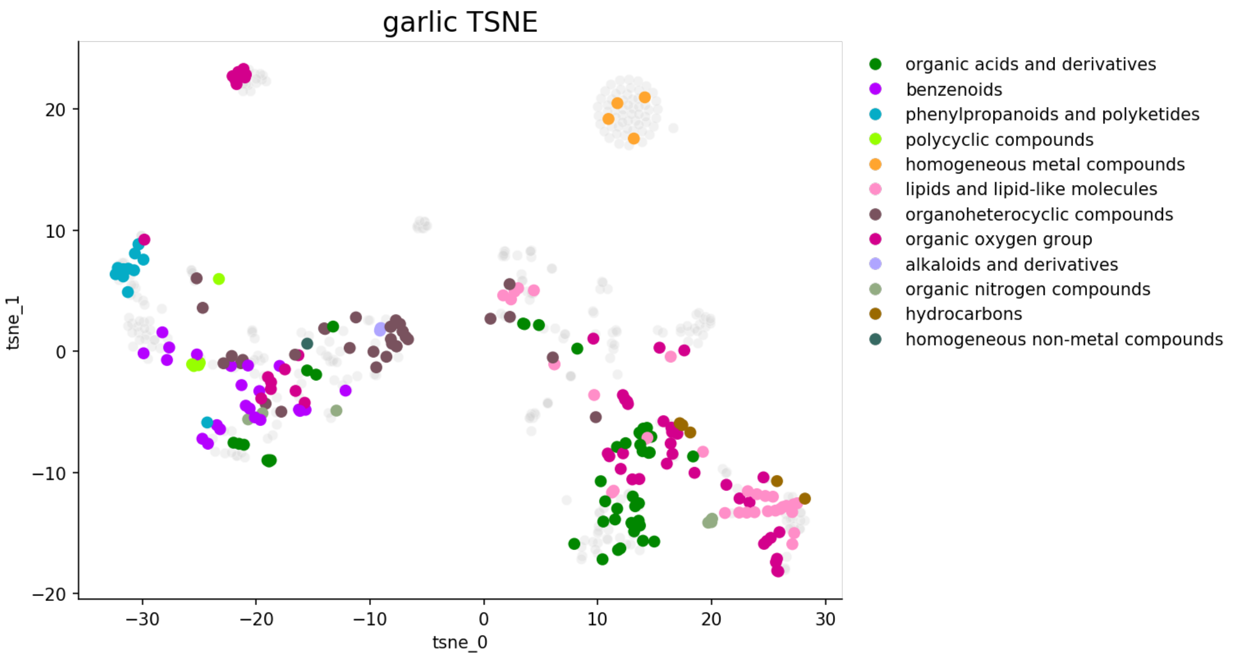


(B)

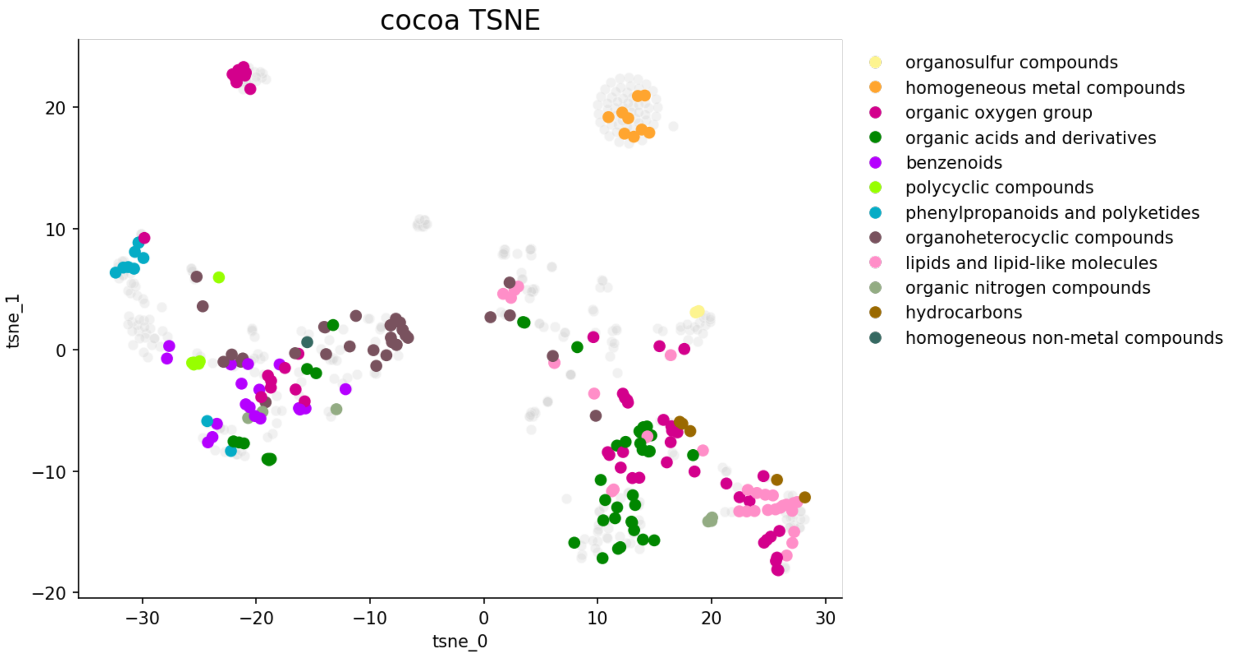


***Figure S5: TSNE Dimensionality Reduction of Chemical Embeddings with Chemical Classes.*** *The plots show the 2-D dimensionality reduction of chemical embeddings for garlic (A) and cocoa (B) chemicals in*

*FoodMine, FooDB, and USDA. We used chemical superclass classifications from FooDB as color code. Grey points represent chemicals that did not have corresponding superclass information.*
